# Quantum entropy reveals chromosomal disorder of ancestry tracts in genetic admixture

**DOI:** 10.1101/2023.02.12.528199

**Authors:** Tianzhu Xiong, Kaifeng Bu

**Affiliations:** Department of Organismic and Evolutionary Biology, Harvard University, Cambridge, MA, USA; Department of Physics, Harvard University, Cambridge, MA, USA

## Abstract

Ancestry tracts are contiguous haplotype blocks inherited from distinct groups of common ancestors. The genomic distribution of ancestry tracts (or local ancestry) provides rich information about evolutionary mechanisms shaping the genetic composition of hybrids. The correlation structure of ancestry tracts has been particularly useful in both empirical and theoretical studies, but there is a lack of *descriptive* measures operating on arbitrarily large genomic blocks to summarize this correlation structure without imposing too many assumptions about admixture. We here develop an approach inspired by quantum information theory to quantify this correlation structure. The key innovation is to represent local ancestry as quantum states, where less correlation in local ancestry leads to elevated quantum entropy. By leveraging a variety of entropy measures on local ancestry signals, we show that entropy is deeply connected to co-ancestry probabilities between and within haplotypes, so that ancestral recombination graphs become pivotal to the study of entropy dynamics in admixture. We use this approach to characterize a standard neutral admixture model with an arbitrary number of sources, and recover entropic laws governing the dynamics of ancestry tracts under recombination and genetic drift, which resembles the second law of thermodynamics. In application, entropy is well-defined on arbitrarily large genomic blocks with either phased or unphased local ancestry, and is insensitive to a small amount of noise. These properties are superior to simple statistics on ancestry tracts such as tract length and junction density. Finally, we construct an entropic index reflecting the degree of intermixing among ancestry tracts over a chromosomal block. This index confirms that the Z chromosome in a previously studied butterfly hybrid zone has the least potential of ancestry mixing, thus conforming to the “large-X/Z” effect in speciation. Together, we show that quantum entropy provides a useful framework for studying ancestry tract dynamics in both theories and real systems.

## 1. Introduction

Closely related species are prone to hybridization in sympatry (Taylor and Larson, 2019). At the genome level, hybridization causes admixture, a process of mixing genomes and eroding species-specific genetic information (Moran et al., 2021). Admixture affects evolution profoundly: it distorts local gene genealogies (Suvorov et al., 2022) and brings distinct mutations/alleles into different species (Skov et al., 2020); it triggers natural selection acting on introgressed genetic elements (Oziolor et al., 2019); this selection process can subsequently strengthen or weaken species boundaries and shape biodiversity (Kleindorfer et al., 2014; Ungerer et al., 1998).

To describe and understand admixture, classical population genetics offer two formalisms. Firstly, properties localized to narrow regions of the genome (“loci”) are studied using measures such as allele frequency divergence (Wright, 1949), linkage disequilibrium (Lewontin and Kojima, 1960), genetic diversity (Nei, 1973), and geographic clines (Barton and Hewitt, 1985). These approaches are mainly descriptive but offers useful insights about the study system (e.g., high *F_ST_* could be a sign of barriers to gene flow, Sakamoto and Innan (2019)). Secondly, one focuses on the entire genome/chromosome and infers haplotype blocks inherited from different ancestral populations. Blocks of constant local ancestry (or “ancestry tracts”) provide correlational information that can be used to study the interplay among recombination, selection, and demography (Pool and Nielsen, 2009; Sedghifar et al., 2016; Shchur et al., 2020; Steinrücken et al., 2018). In practice, analyzing local ancestry often begins with imposing a particular model of admixture, then estimating parameters such as admixture times, selection coefficients, or effective population sizes (Gravel, 2012; Liang et al., 2022; Svedberg et al., 2021), which could be highly complex with many degrees of freedom. Interpreting and comparing these complex models is challenging because evolutionary processes are likely heterogeneous along the genome, and it is often unknown *a priori* if a model is misspecified (Huang et al., 2022).

Given the complexity of admixture at genomic scales, there is currently a lack of frameworks allowing researchers to simply *observe* the correlation structure of ancestry tracts in the study system without making too many assumptions about models. This can be particularly useful when local ancestry is easy to infer (e.g., ancestry-informative SNPs are dense in the genome). Furthermore, from a theoretical perspective, descriptive measures on ancestry tracts will be useful in quantifying evolutionary dynamics with a global impact on genomes or chromosomes, such as coupling among polygenes underlying reproductive isolation (Feder et al., 2014), interactions between assortative mating and hybrid incompatibility (Muralidhar et al., 2022), or large structural variants affecting chromosomal recombination (de Vos et al., 2020; Kirkpatrick and Barton, 2006). Understanding these nonlocal processes requires quantitative measures operating on arbitrarily large genomic blocks.

To date, a few summary statistics have been developed to measure local ancestry correlation, among which ancestry tract length and ancestry junction density are the most widely used (Baird et al., 2003; Fisher, 1949, 1954; Janzen et al., 2018; Liang and Nielsen, 2014). Both measures describe how frequently local ancestry switches states along a haplotype. This information has been used not only in timing admixture events, but also in detecting natural selection during admixture, because selected sites are physically associated with longer ancestry tracts, or equivalently, fewer ancestry junctions — a signature of less mixed local ancestry (Shchur et al., 2020; Wang et al., 2022). Although both measures are intuitive, problems exist in practice. First, tract length requires high-quality haplotypes because length is undefined if ancestry is unphased in diploid organisms. Second, as a counting measure, junction density is sensitive to even localized errors in ancestry inference. Similarly, small errors in local ancestry inference can rupture an otherwise contiguous ancestry tract, producing bias towards shorter tracts (Harris and Nielsen, 2013; Ralph and Coop, 2013). This is particularly problematic if phase imputation is forced in the absence of reliable haplotype information. Lastly, many studies identify a specific ancestry as the major parent, while treating the rest as introgression, hence restricting admixture to only two sources with an inherent polarity in analysis (Schumer et al., 2018; Vilgalys et al., 2022). This could be problematic for samples in the center of a hybrid zone since the major parent might change along the genome when contributions are relatively equal among all sources.

Inspired by quantum information theory, here we develop a descriptive framework to quantify ancestry mixing in an arbitrarily large genomic block with any number of source populations. It has clear population genetic interpretations with phased local ancestry, and it is adjustable for unphased data. Based on a prototype, the core of this framework is the entropy over a set of local ancestry signals (Xiong et al., 2022a). We will start by representing genetic ancestry as quantum states. Then we will define entropy and connect it to co-ancestry probabilities that are inferrable from ancestral recombination graphs, thus grounding the current framework in coalescent theory. Finally, we show the usefulness of entropy as descriptive statistics to study admixture dynamics at genomic scales.

## 2. Results

### 2.1 Intuition for the quantum representation of genetic ancestry

An obstacle in the mathematical representation of genetic ancestry is that “ancestry” is an unordered categorical variable. For instance, modern humans inherit multiple types of ancestry from ancient populations (Gopalan et al., 2022), but ancestral human populations cannot be ordered into sequential relationships. This implies that a mathematical representation of ancestry must be multidimensional and tracks contributions from all sources simultaneously. In quantum physics, information is encoded in quantum states, which provides a probability distribution of all possible measurement outcomes (Nielsen and Chuang, 2010). These outcomes are usually categorical and unordered, such as the spin of particles. In application, the correlation structure of genomic ancestry tracts provides invaluable information about evolutionary mechanisms, while the correlation structure of quantum states is also essential for the study of quantum entanglement (Horodecki et al., 2009) and the related quantum information theory. This resemblance between genetic ancestry and quantum states inspires us to borrow the existing framework in quantum information theory to study genetic admixture. Nonetheless, nothing is fundamentally quantum in population genetics, and our approach should be understood as a formal appropriation of concepts and methods in quantum information theory. A quantum state is usually denoted as |·〉, an element in a complete vector space equipped with an inner product. Its adjoint vector is written as 〈·|, and 〈·|·〉 represents inner products. In addition, averaging a function over all of its positional arguments *l*_1_, *l*_2_,… is written as 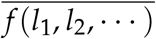 (SI Definition 1).

To represent genetic ancestry, let |*k*〉 be a pure state corresponding to the *k*-th source population (or the *k*-th ancestry type). As many organisms have ploidies larger than one, measuring ancestry using unphased genotypes will produce mixed ancestry at a single locus. We propose that both pure and mixed local ancestries can be represented by the superposition of pure states (Fig. 1A):

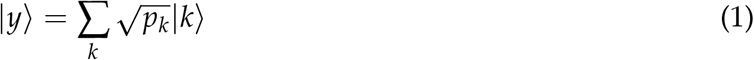

where *p_k_* corresponds to the contribution from the *k*-th source. Naturally, ∑_*k*_ *P_k_* = 1. For instance, heterozygous ancestry in a diploid individual with two admixture sources is

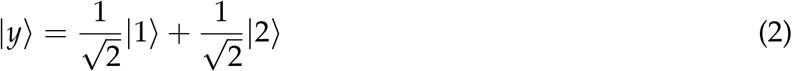

**Figure 1:**
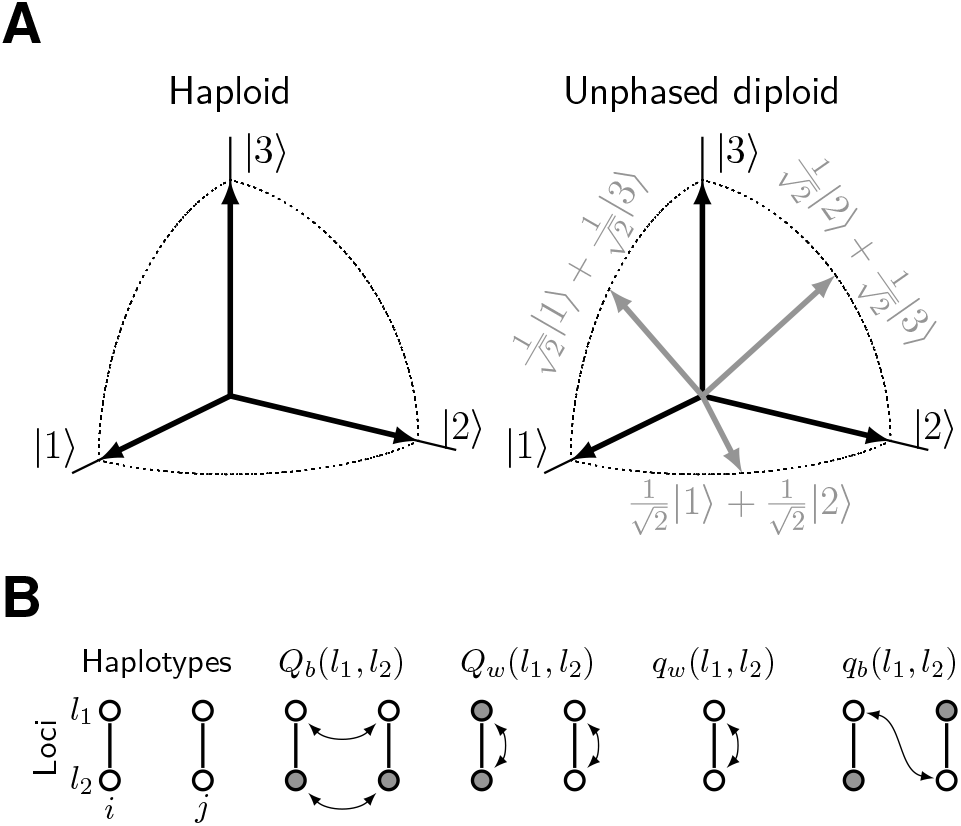
Key elements of the framework. **(A)** An example of the quantum representation of genetic ancestry for individuals with three orthogonal sources of admixture. For diploids, gray arrows are loci heterozygous in ancestry, and black arrows are loci homozygous in ancestry. **(B)** Probabilities of co-ancestry. These probabilities are defined on ancestries at two positions (*l*_1_ and *l*_2_) on one or two haplotypes (*i* and *j*). Alleles connected by arrows are identical by ancestry.

Each source population is often treated as a distinct entity. Thus, pure states are assumed to be orthogonal to each other:

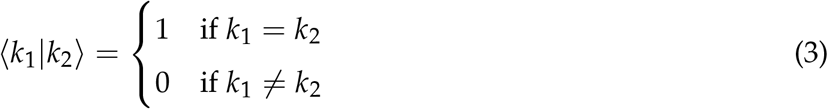

Orthogonality also enables a probabilistic interpretation of entropy to be introduced later.

In practice, local ancestry is the outcome of inheritance history, and it needs to be inferred from genome sequences with varying levels of differentiation. If parental populations are not differentiated at the sequence level, ancestry is not observable, but the validity of our theory is not affected since inheritance history always exists^1^.

### 2.2 Entropy associated with ancestry correlation over chromosomal blocks

There are two major axes of ancestry correlation: the one between chromosomal positions and the one between individuals. For a contiguous chromosomal block with positional index *l* ∈ [0, *L*], suppose a sample contains *n* individuals (*n* ancestry signals). Define the following continuous integral kernel:

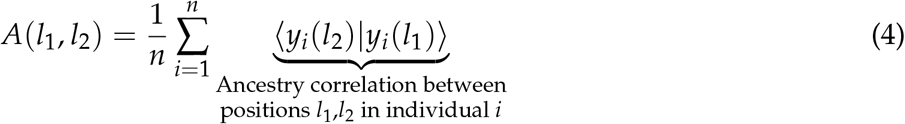

Kernel *A* captures pairwise ancestry correlation between chromosomal positions and averaged over all individuals. Next, define the following kernel matrix **B** = {*b_ij_*}_*n×n*_, where

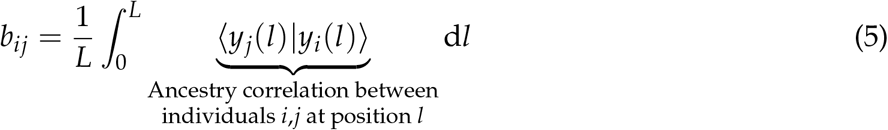

Matrix **B** captures pairwise ancestry correlation between individuals and averaged over all positions in the chromosome block. Each kernel has a unique spectral decomposition with real eigenvalues:

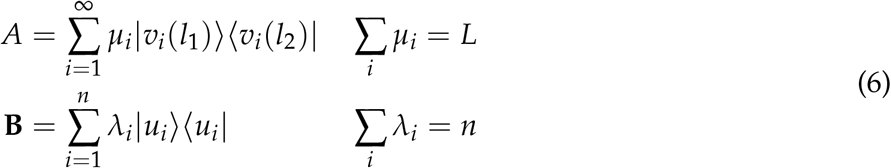

For these kernels, less amount of correlation between different components increases the evenness of eigenspectrum, and this evenness can be measured by quantum entropy *S*. Two common forms of quantum entropy are the so-called “von Neumann entropy” and “linear entropy” (Eq. 7). Widely used in physics, von Neumann entropy is simply the classical Shannon entropy formula applied to the kernel spectrum, but it is an extensive quantity and grows with *O*(log *n*). On the contrary, linear entropy is nonextensive and is always bounded by one. To overcome sample size dependency, we adopt linear entropy for all subsequent analyses, and entropy will be used synonymously with linear entropy unless explicitly specified.

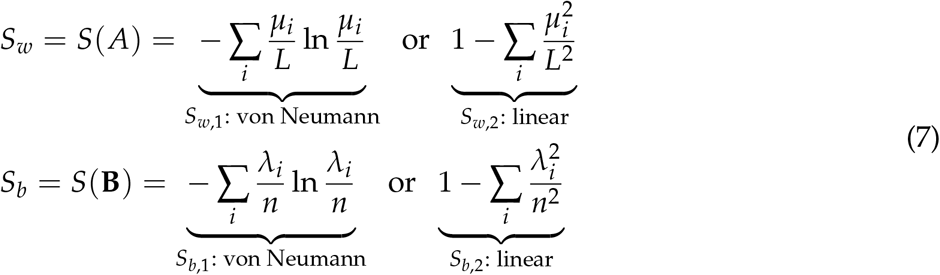

The subscript “*w*” stands for “within-individual”, and “*b*” stands for “between-individual”, which are understandable from the axes of correlation. Note that an equivalent expression for linear entropy is to use the trace operator “Tr”:

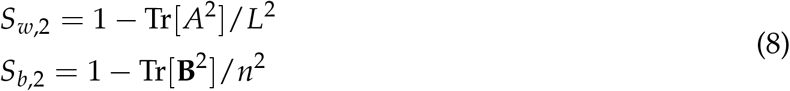

### 2.3 Linear entropy captures co-ancestry probabilities among haplotypes

A nice property of linear entropy is that it has a clear population genetic interpretation when samples are haplotypes. For *n* haplotypes, consider the following probabilities of co-ancestry (Fig. 1B): Let *Q_b_*(*l*_1_, *l*_2_) be the probability that two different haplotypes share ancestry at both loci *l*_1_ and *l*_2_, and let *Q_w_* (*l*_1_, *l*_2_) be the probability that two loci *l*_1_ and *l*_2_ share ancestry in both haplotypes. Further, let *q_w_* (*l*_1_, *l*_2_) be the probability that two loci share ancestry in a single haplotype.

The expectation of linear entropy is related to these probabilities of co-ancestry by (SI Theorem 1):

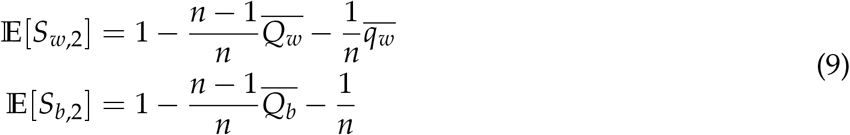

where 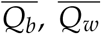, and 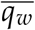 are the corresponding probabilities averaged over all pairs of loci in the chromosome block [0, *L*]. One can adjust sample size *n* to select information most interesting to the problem. If correlation along individual haplotypes is the only information to consider, take *n* = 1 and this information is encoded in *S*_*w*,2_ as

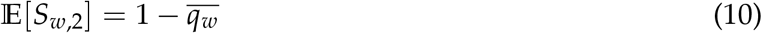

Here, 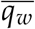 is simply the two-point correlation function of local ancestry averaged over a chromosomal block, and it is closely related to linkage disequilibrium in classical population genetics. In the limit of large sample sizes, more complex correlation information is encoded in entropy:

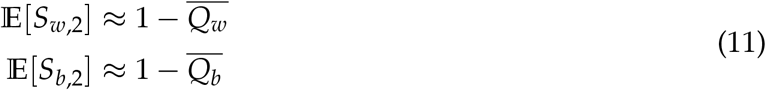

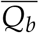 and 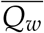 contain more correlation information than linkage disequilibrium because they are also affected by ancestry correlation between haplotypes (Fig. 1B), thus incorporating the disorderedness generated by both recombination and genetic drift.

### 2.4 A standard model: entropy dynamics among haplotypes in neutral admixture

As a proof of concept, we apply entropy to a simple but standard admixture model to test if it can be used to track ancestry disorderedness. In this model, *n* haplotypes are sampled from a large, diploid, and randomly mating population of size *N*. The population is established by admixture at time *t* = 0 from *K* sources with proportions **p** = (*p*_1_,…, *p_K_*), and it evolves neutrally under the Wright-Fisher process with identical recombination rates at all chromosomal positions with no interference (Durrett, 2008). This multi-source neutral admixture is a useful approximation for closely related populations. A single admixture event is assumed to demonstrate the fundamental properties of entropy.

The probabilistic interpretation of linear entropy enables us to investigate its dynamics analytically (see SI “Ancestral recombination graph”). Overall, the expected entropy changes on two timescales (top parts in Fig. 2A): recombination within blocks increases entropy with a rate determined by block-average recombination probabilities, and genetic drift dampens entropy with a rate determined by effective population sizes. When the two timescales differ greatly (e.g., long blocks in a large population), a chromosomal block will experience a rapid burst of entropy, followed by a slow decline due to genetic drift removing diverse haplotypes. This behavior is accurately captured by forward-in-time simulations (colored curves in Fig. 2B) and analytical models using ancestral recombination graphs (dashed and dotted curves in Fig. 2B).

**Figure 2:**
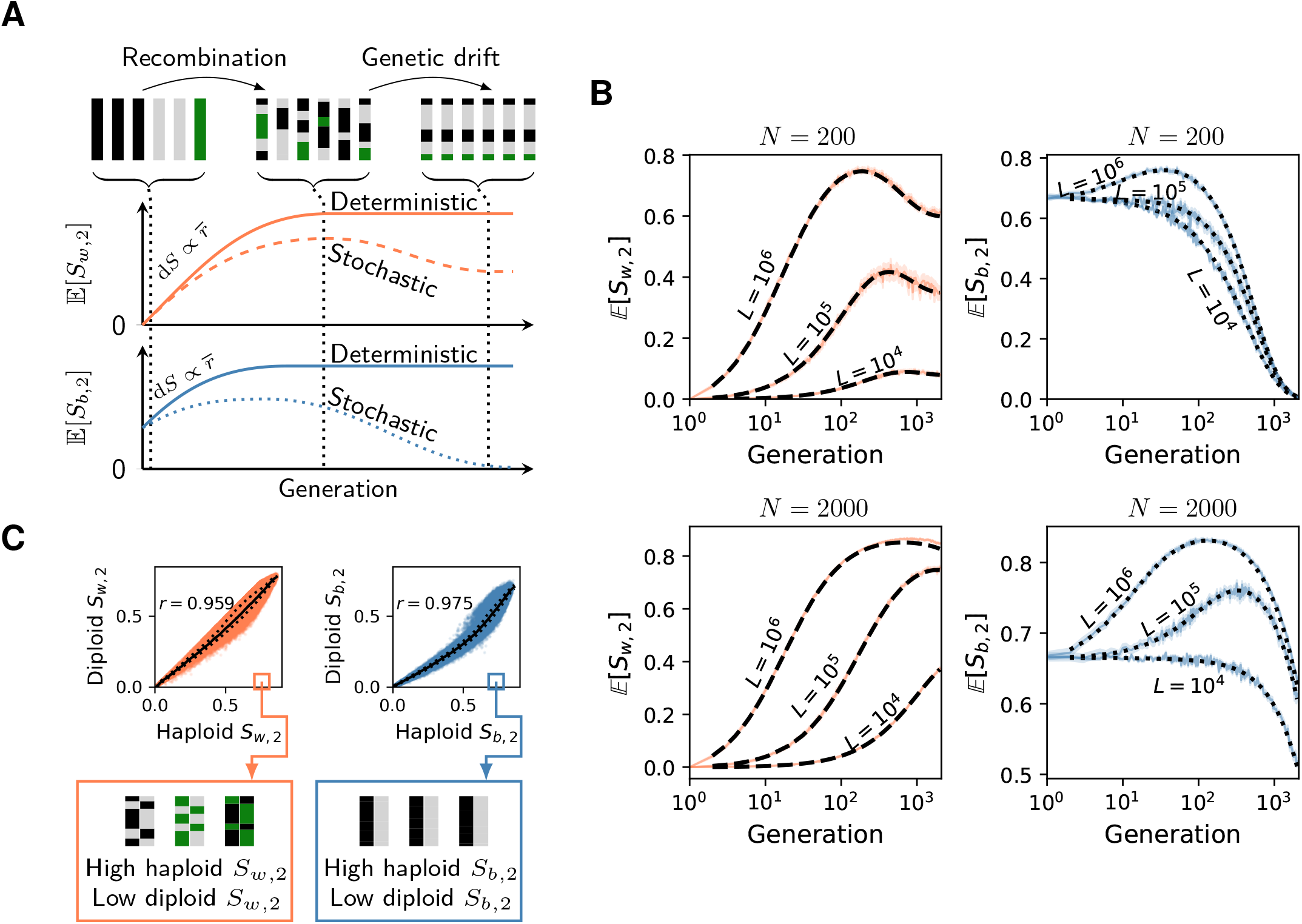
Entropy dynamics among haplotypes in neutral admixture. **(A)** A long chromosomal block will first accumulate entropy rapidly due to recombination breaking up ancestry correlation within and between haplotypes, but stochasticity such as genetic drift will eventually dampen entropy to a lower equilibrium. Curves are illustrative and are not based on exact numerical values. **(B)** Simulated (with SLiM-4.0) and analytical trajectories of the expected entropy. For simulation, each individual has a pair of homologous chromosomes of length 10^7^ bp. Recombination occurs independently between adjacent sites with a probability 10^−7^ per generation. Admixture proportion **p** = (0.1, 0.2, 0.3, 0.4). The expected entropy is computed for 10 diploid individuals (*n* = 20) by averaging entropy among non-overlapping chromosomal blocks of length *L*, then averaging over multiple permutations of individual ancestry assignment at *t* = 0, and finally averaging over five independent simulations of the whole model. Colored boundaries are mean +/− standard error of the expected entropy. Dashed and dotted curves are the expected entropy evaluated using ancestral recombination graphs. **(C)** Top: Relationships between haploid and diploid entropy measures in neutral admixture. Each dot represents a long block (*L* = 10^6^ bp) in an admixed population (*N* = 200) in the aforementioned simulation with the same admixture proportion. Black solid lines and dotted lines are median and (25%, 75%) percentiles of diploid entropy, respectively. *r* is Pearson’s correlation coefficient. Bottom: Some ancestry configurations are associated with diploid entropy significantly underestimating haploid entropy.

#### Short-term behavior

Let *r*(*l*_1_, *l*_2_) be the recombination probability per generation between loci *l*_1_ and *l*_2_. Ancestral recombination graphs predict the following rates for the short-term production of entropy at the onset of admixture (SI Theorem 4):

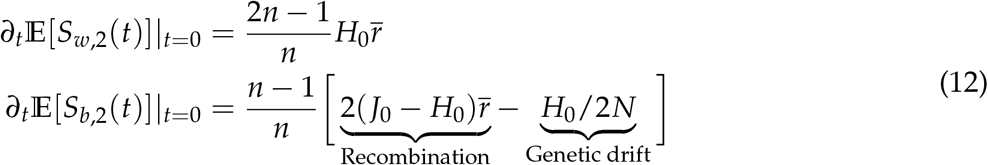

where 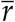 represents recombination probabilities averaged over all pairs of loci in a chromosomal block, and (*H*_0_, *J*_0_) are two constants determined by admixture proportions:

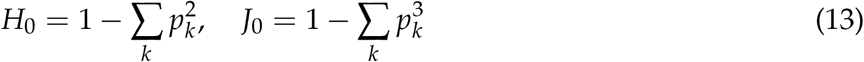

Eq. 12 shows that the initial production of both entropies is proportional to the block-average re-combination probability 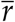. The dependency of entropy production on 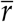 has two implications: when per-base-pair recombination rate is fixed, longer blocks will on average accumulate entropy faster (Fig. 2B); conversely, if block size is fixed, recombination hotspots will on average accumulate entropy faster than regions with suppressed recombination. The effect of genetic drift will be prominent if a chromosomal block is sufficiently short so that

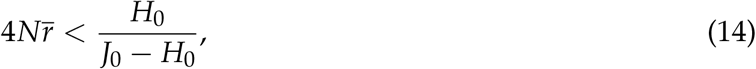

causing the immediate decrease of 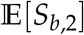 at the onset of admixture.

#### Long-term behavior

Without genetic drift, recombination elevates the expected entropy to the following deterministic limits (SI Theorem 5):

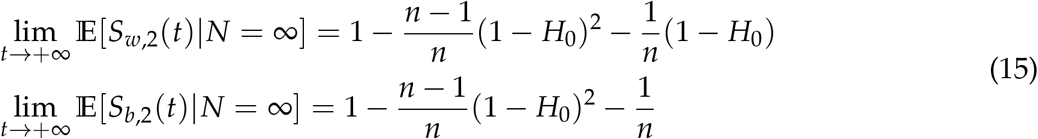

Due to genetic drift, these limits are rarely attained in a small population or on a very short chromosomal block. Eventually, the loss of haplotype diversity dampens entropy to these stochastic long-term limits (SI Theorem 6):

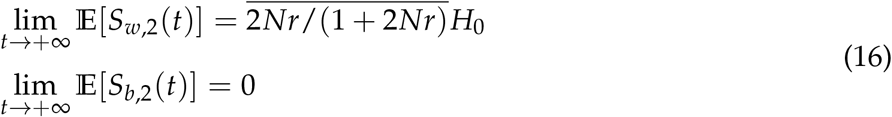

This sets apart the dynamics of entropy in admixed but highly inbred populations from those in populations with very large effective population sizes (compare deterministic and stochastic trajectories in Fig. 2A, and between top/bottom panels in Fig. 2B). In particular, long-term equilibria of 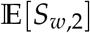 decreases with a smaller *N*, implying that increased genetic drift fixes longer ancestry tracts, a result congruent with classical studies of ancestry tract lengths(Gravel, 2012). We find that the expected total entropy in this standard model satisfies the following inequality, which we call the “**Heisenberg uncertainty principle for neutral genetic admixture**”^2^:

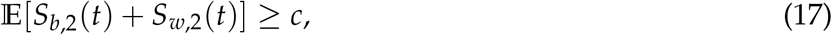

where 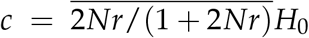 characterizes the competition between recombination and genetic drift.

#### Diploid vs haploid entropy

While entropy among haplotypes is a natural measure of the disorderedness of ancestry tracts, reliable haplotypes are costly to assemble, and local ancestry is often unphased in empirical studies. The diploid ancestry representation (Fig. 1A) relaxes the phasing requirement and entropy can be evaluated on *n* unphased diplotypes. Does diploid *S* capture information similar to haploid *S*? In simulated neutral admixture, the two measures are strongly correlated (Fig. 2C: top), and diploid *S* is always the smaller one. For them to differ greatly, diploid *S* must severely underestimate haploid *S*. For *S*_*w*,2_, this can happen only when diploid ancestry is mostly heterozygous while haplotypes contain highly broken ancestry tracts (Fig. 2C: bottom left). However, this scenario is statistically unlikely because it requires local ancestry between the two haplotypes to be anti-correlated and ancestry junctions to be aligned. In a large population, ancestry junctions are unlikely to be shared between haplotypes; and even if they are shared via identical-by-descent, local ancestry should be the same rather than be anti-correlated. Consequently, diploid *S*_*w*,2_ is still a good measure in the absence of phase information. For diploid *S*_*b*,2_, its behavior can be less reliable. For instance, in a sample consisted entirely of F1 hybrids, haploid *S*_*b*,2_ is large, but diploid *S*_*b*,2_ becomes zero because all individuals have the same genotype (Fig. 2C: bottom right). This problem occurs because F_1_ hybrids are far from Hardy-Weinberg equilibrium. In subsequent generations this problem becomes less severe and diploid *S*_*b*,2_ is still strongly correlated with haploid haploid *S*_*b*,2_.

### 2.5 Irreversible change in ancestry tracts resembles the second law of thermody-namics

In thermodynamics, entropy change is often associated with irreversible processes (the second law of thermodynamics, Sethna (2021)). In population genetics, ancestry tracts broken up by recombination cannot be systematically stitched back by chance. Similarly, haplotypes lost to genetic drift cannot spontaneously reappear in the absence of other processes. This directionality is visible from Fig. 2A, that these neutral processes shift ancestry correlation from *within* to *between* haplotypes. Pictorially, in the genotypic space of all possible haploid ancestry signals, an admixed population will spontaneously evolve from haplotypes forming multiple clusters to a single one. To capture this intrinsic directionality in admixture dynamics, we introduce:

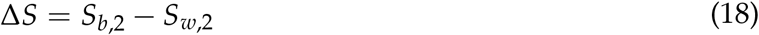

With large sample sizes, 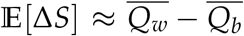, which is exactly the difference between the two probabilities of co-ancestry in Fig. 1B. Thus, the sign of Δ*S* tells the major axis of co-ancestry (within vs between), and a significantly positive Δ*S* coincides with the presence of distinct ancestry clusters in that genotypic space.

To illustrate this property, we track Δ*S* in the neutral admixture model through time (Fig. 3). The initial generation has haploid Δ*S* = *H*_0_ > 0, congruent with the presence of distinct multilocus ancestry clusters (Fig. 3: *t* = 1). In an infinitely large population, recombination induces an irreversible change in haploid Δ*S* (SI Theorem 7):

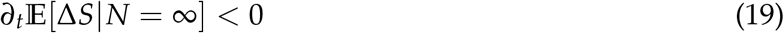

**Figure 3:**
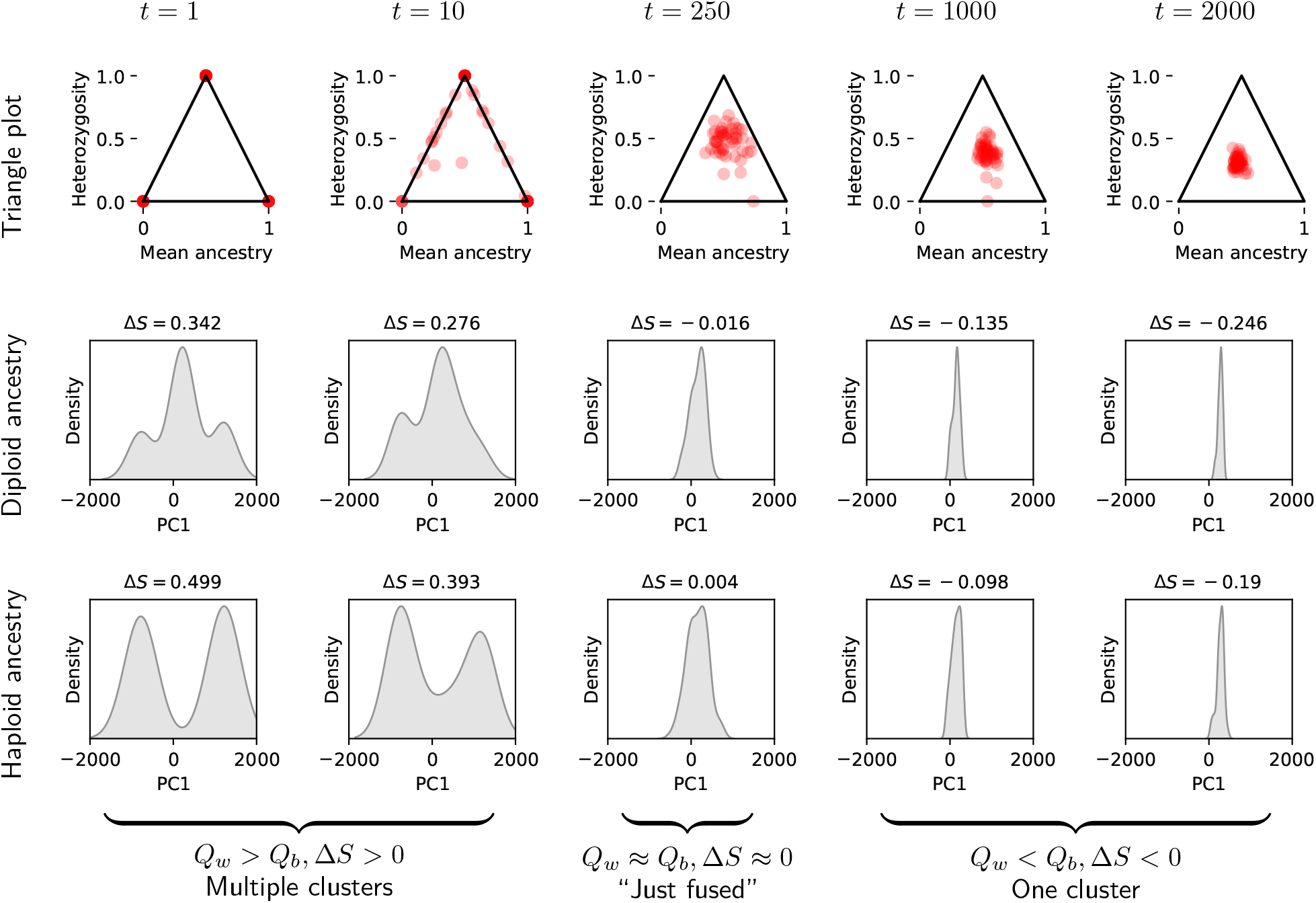
Entropy change reveals the fusion and contraction of clusters in genotypic space. For each *t* (generation), 50 diploid individuals are sampled from a population (*N* = 2000) with twoway admixture (**p** = (0.5, 0.5)). All quantities are based on local ancestries of a single chromosomal block (*L* = 10^6^ bp) across sampled individuals. Genotypic clustering is visualized using principal component analysis on local ancestry. The first principal component (PC1) is computed from the first-generation sample and all later-generation samples are projected onto the same PC1. All principal component analyses use numerically transformed ancestry as follows: haploids with two states: (1, −1); and diploids with three states: (1,0, −1). Only the density along PC1 is shown for each distribution (fitted using a Gaussian kernel). **Top row:** Each dot represents a diploid individual’s mean ancestry and heterozygosity of a given block. **Middle row:** The sign of Δ*S* using diploid entropy is correlated with the distribution of individuals along the first principal component (PC1) of diploid ancestry. **Bottom row:** The sign of Δ*S* using haploid entropy is correlated with the distribution of haplotypes along the first principal component (PC1) of haploid ancestry.

This shows that recombination causes ancestry clusters to fuse. Haploid Δ*S* in the final stage of deterministic admixture is close to zero (Fig. 3: *t* = 100):

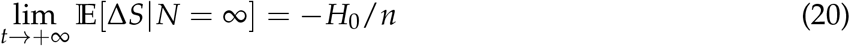

This occurs when ancestry tracts have fully disintegrated and co-ancestry probabilities between and within haplotypes are similar. In finite populations, co-ancestry probability *Q_b_* will gradually rise between haplotypes due to genetic drift, and Δ*S* will decrease further (Fig. 3: *t* = 2000). We use numerical methods to show that 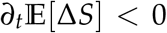 is valid under both genetic drift and recombination (SI Conjecture 1, Fig. S2 and S3). For sufficiently long blocks, the stochastic long-term limit of 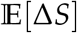 is precisely the opposite of the initial entropy (*H*_0_), which implies that

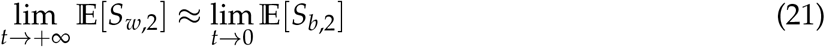

This relation shows that, among long haplotype blocks, recombination and genetic drift transfer approximately all the initial entropy *between* haplotypes in the first generation to the entropy *within* a single haplotype at equilibrium. The change in diploid Δ*S* is similar (middle row in Fig. 3).

### 2.6 A chromosomal measure of admixture progress based on entropy diagrams

Ancestry signals are high-dimensional, and distilling their correlation structure into entropy enables the use of low-dimensional diagrams for qualitative studies. Information provided jointly by *S*_*b*,2_ and *S*_*w*,2_ can already separate different kinds of ancestry signals (Fig. 4A), and Δ*S* splits them by the presence of distinct ancestry clusters. Nonetheless, similar to linkage disequilibrium in classical population genetics, the magnitude of Δ*S* is affected by the frequency of each source. For instance, Δ*S* is zero in a pure population without admixture as well as in a population with fully disassociated ancestry tracts (the diagonal line in Fig. 4A). While these populations contain a single ancestry cluster, they are “homogeneous” with very different levels of ancestry variation. Thus, Δ*S* alone is insufficient to probe the degree of intermixing among ancestry tracts.

**Figure 4:**
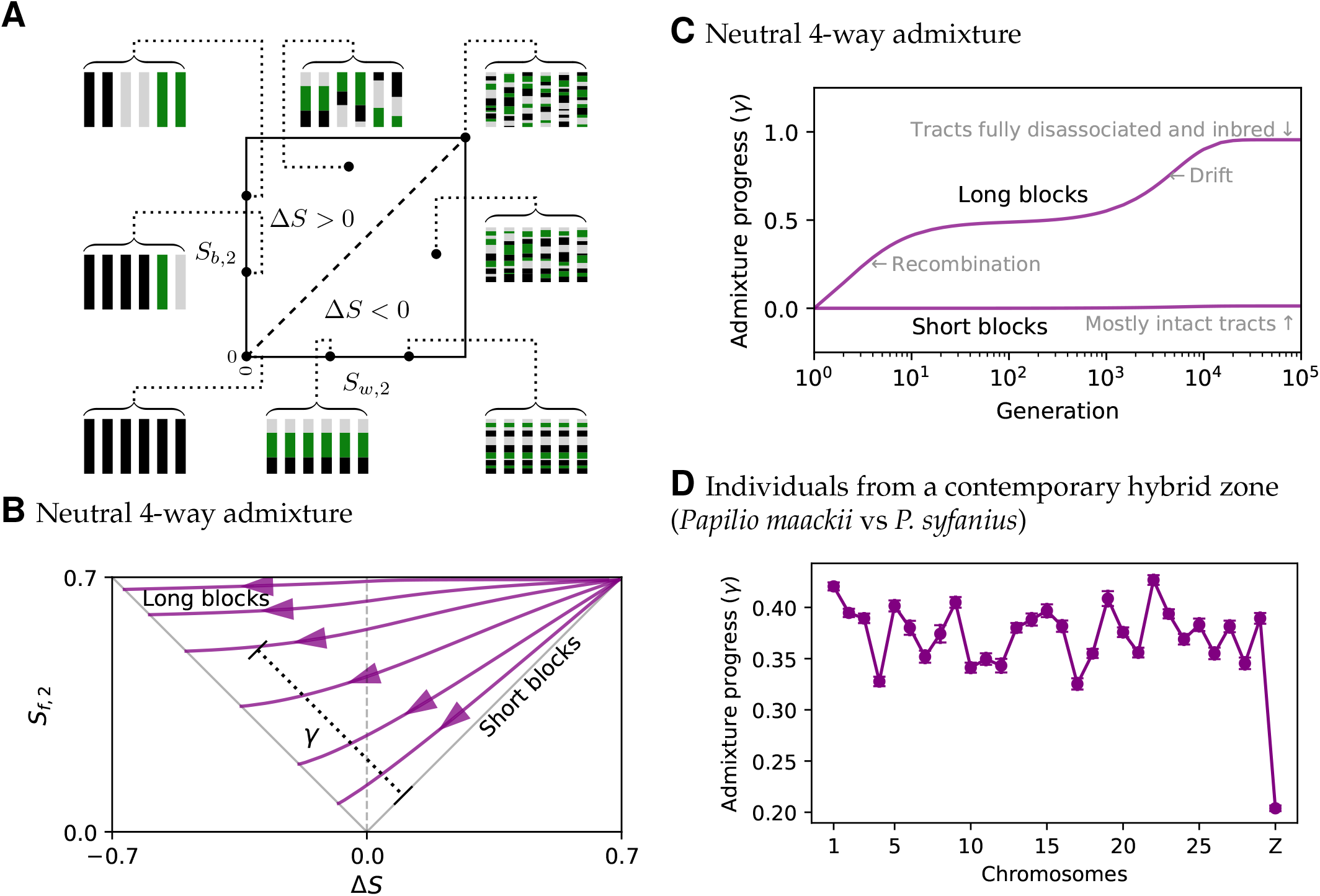
Entropy diagrams. **(A)**. The diagram between *S*_*b*,2_ and *S*_*w*,2_ visualizes correlation structures of ancestry signals. The upper-left triangle has Δ*S* > 0, and ancestry is more correlated within haplotypes. The lower-right triangle has Δ*S* < 0, and ancestry is more correlated between haplotypes. Schematic haplotypes with three sources are shown for selected points in the diagram. **(B)**. The diagram between Δ*S* and *S*_*f*,2_. Purple curves are the expected trajectories computed for neutral admixture among haplotypes using ancestral recombination graphs. Model parameters: **p** = (0.1, 0.2, 0.3, 0.4), *n* = 100, *N* = 2000, *L* ∈ {10^3^bp,3162bp, 10^4^bp, 31623bp, 10^5^bp,10^6^bp}. Recombination probability= 10^−7^ per site per generation. Measure *γ* assesses admixture progress by calculating the distance to line *S*_*f*,2_ = Δ*S* in the diagram (i.e., the trajectory taken by an extremely short block). **(C)**. Short blocks maintain tract integrity for many more generations in neutral admixture due to a lack of recombination. Curves are computed for the same model as in the previous figure. *L* ∈ {10^2^bp, 10^7^bp}. **(D)**. *γ* computed for each chromosome from a contemporary butterfly hybrid zone shows a significant reduction of admixture progress on the sex chromosome (Z chromosome).

As a remedy, we augment Δ*S* by the total amount of ancestry variation for a given block across all sampled individuals. This can be quantified by the entropy of the fractions (**F**) of different sources.

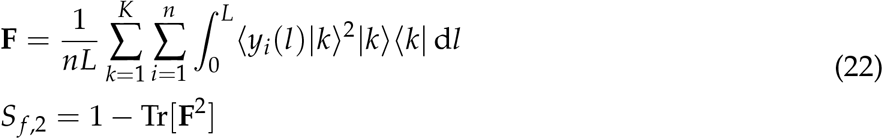

Similarly, *S*_*f*,2_ has a probabilistic interpretation (SI Theorem 2):

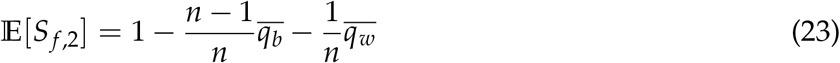

where *q_b_*(*l*_1_, *l*_2_) is the probability that position *l*_1_ in one haplotype shares ancestry with position *l*_2_ in a different haplotype (Fig. 1B). *S*_*f*,2_ is independent of phasing since **F** erases haplotype and individual information.

Together, signatures of ancestry clusters (Δ*S*) and the total amount of ancestry variation (*S*_*f*,2_) can be used to visualize ancestry dynamics. For instance, a general feature of neutral admixture is that short blocks maintain distinct ancestry clusters more easily than longer blocks because the number of recombination events scales with block size. This feature is captured by a simple diagram between *S*_*f*,2_ and Δ*S* (Fig. 4B): All blocks start from a high level of ancestry variation with distinct clusters (top-right of Fig. 4B); For long blocks, ancestry tracts quickly mix up by recombination, so *S*_*f*,2_ is nearly a constant while Δ*S* decreases (the top trajectory in Fig. 4B). For short blocks, their low recombination rates allow genetic drift to dampen *S*_*f*,2_ and Δ*S* simultaneously (the lower-right trajectory in Fig. 4B). Since the trajectory taken by an extremely short block represents no intermixing among ancestry tracts, a natural measure to assess admixture progress is to compute the distance (*γ*) between a point in this diagram and the trajectory taken by very short blocks (the line Δ*S* = *S*_*f*,2_). As expected, *γ* on long blocks increases rapidly during neutral admixture, while *γ* on short blocks stays close to zero (Fig. 4C).

As an example in real hybrids, we computed *γ* for each chromosome from a butterfly hybrid zone, which has already been characterized by a preliminary version of this entropy approach (Xiong et al., 2022a). Here, we mapped reads to a newly corrected reference genome and inferred unphased local ancestry on four individuals, demonstrating that the sex chromosome (Z chromosome) has much less admixture in the hybrid zone (Fig. 4D). This corroborates the observation that the Z chromosome is either pure in ancestry or with ancestry tracts resembling those in a backcross individual (Fig. S4).

### 2.7 A chromosomal measure of parallel evolution between hybrid populations

To this point, we have limited our analysis to one sample from a single hybrid population. When multiple hybrid populations exist, a natural question is whether admixture proceeds predictably between different hybrid zones so that they share the same underlying selection process. In terms of ancestry signals, it is equivalent to asking if ancestry varies in similar ways across the genome between samples from different populations. Suppose two samples of sizes *n*_1_ and *n*_2_ are taken from two hybrid populations. The pairwise ancestry correlation between individuals from different populations is captured by

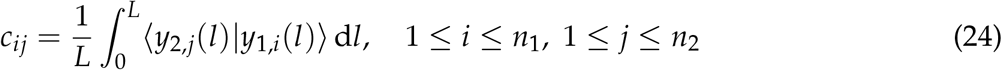

where |*y*_1,*i*_〉 represents the local ancestry of individual *i* from the first population (|*y*_2,*j*_〉 is interpreted similarly). These elements form an *n*_1_ × *n*_2_ dimensional matrix **C** with a corresponding entropy

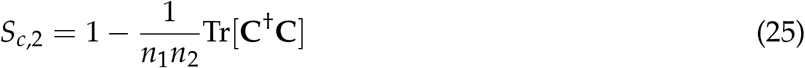

where **C**^†^ is the conjugate transpose of **C**. The subscript “*c*” stands for “cross-population”. The probabilistic interpretation of this entropy is simply (SI Theorem 3):

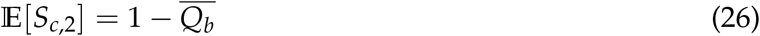

where *Q_b_*(*l*_1_, *l*_2_) is interpreted as the probability that two haplotypes, one from each population, share ancestry at both loci *l*_1_ and *l*_2_ (see Fig. 1B). This probability increases when different hybrid populations fix ancestry in a concerted way along the genome. Thus, highly predictable and parallel evolution between different populations will be associated with a lower *S*_*c*,2_.

## 3. Discussion

### 3.1 Summary of the framework

A major aim of this framework is to provide quantitative, descriptive measures on local ancestry that can be used to summarize features of ancestry tracts. This framework goes beyond simple counting measures such as tract length and junction density, and each different axis of disorderedness in local ancestry can be measured by its corresponding entropy (Table 1). These entropies are associated with several derived measures. First, we showed that the change in Δ*S* = *S*_*b*,2_ − *S*_*w*,2_ reflects the irreversibility of ancestry tract dynamics under recombination and genetic drift. The sign of Δ*S* can also be used to detect the major axis of ancestry correlation. Second, we combined Δ*S* and *S*_*f*,2_ in a diagram and showed that the degree of intermixing among ancestry tracts can be observed directly. By calculating “admixture progress” *γ* in this diagram, the degree of intermixing becomes measurable for arbitrarily large genomic blocks. Finally, we showed that *S*_*c*,2_ can be used to quantify parallel evolution between different hybrid populations, enabling the detection of genomic regions that evolve under similar selection pressures.

**Table 1:**
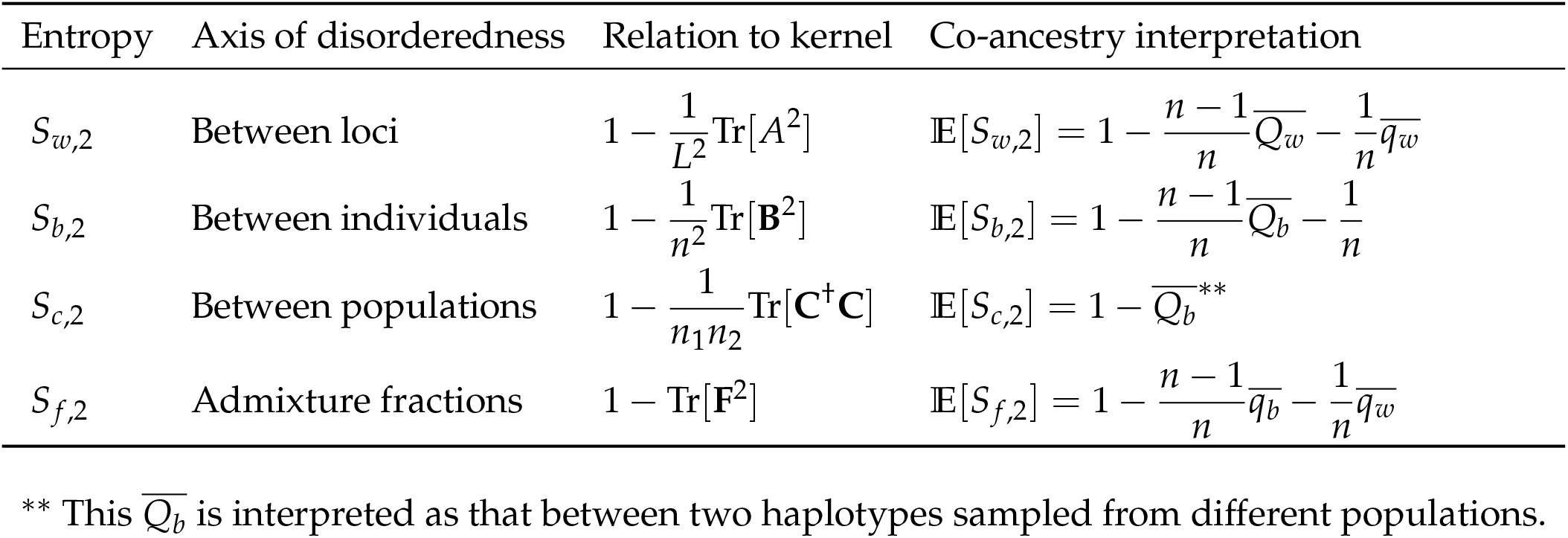
Overview of entropy measures on local ancestry signals

| Entropy | Axis of disorderedness | Relation to kernel | Co-ancestry interpretation |
| --- | --- | --- | --- |
| $S_{w,2}$ | Between loci | $1 - \frac{1}{L^2} \text{Tr}[A^2]$ | $\mathbb{E}[S_{w,2}] = 1 - \frac{n-1}{n} \overline{Q_w} - \frac{1}{n} \overline{q_w}$ |
| $S_{b,2}$ | Between individuals | $1 - \frac{1}{n^2} \text{Tr}[\mathbf{B}^2]$ | $\mathbb{E}[S_{b,2}] = 1 - \frac{n-1}{n} \overline{Q_b} - \frac{1}{n}$ |
| $S_{c,2}$ | Between populations | $1 - \frac{1}{n_1 n_2} \text{Tr}[\mathbf{C}^\dagger \mathbf{C}]$ | $\mathbb{E}[S_{c,2}] = 1 - \overline{Q_b}^{**}$ |
| $S_{f,2}$ | Admixture fractions | $1 - \text{Tr}[\mathbf{F}^2]$ | $\mathbb{E}[S_{f,2}] = 1 - \frac{n-1}{n} \overline{q_b} - \frac{1}{n} \overline{q_w}$ |

\*\* This $\overline{Q_b}$ is interpreted as that between two haplotypes sampled from different populations.

### 3.2 Robustness to erroneous ancestry inference

A key feature of entropy, or similar quantities based on the eigenspectrum of correlation kernel is that each locus contributes evenly to the final result. If a small proportion *ϵ* of ancestry is wrongly inferred, its impact on entropy will also be on the order of 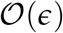, because (linear) entropy is a polynomial function of all kernel elements. On the other hand, this level of error will unpredictably affect ancestry tract length and junction density. There is no theoretical limit to the number of junctions one could falsely pack in a genomic interval of length *Lϵ*. Similarly, for a long ancestry tract, an infinitely small block of wrongly inferred ancestry inside the tract will drastically change tract length. Thus, entropy is more stable to small errors in ancestry inference, and its accuracy depends continuously on error proportion *ϵ*.

### 3.3 Estimation problem

Our framework does not estimate entropy when ancestry is a hidden variable not directly observed. If local ancestry is already known (e.g., inferred with high confidence using HMM models. See Wu et al. (2021) for a comprehensive survey), the framework can be directly applied to either phased or unphased signals. Since all entropy measures have co-ancestry interpretations when samples are haplotypes, many existing tools using ancestral recombination graphs can be used in estimating entropy from haplotype sequences. A future research question is to estimate entropy directly from sequence data while integrating over all hidden local ancestry states.

### 3.4 More complex correlation structures

Linear entropy is a special case (*q* = 2) of the more general Tsallis entropy of a (normalized) correlation kernel *ρ* (Furuichi et al., 2004; Tsallis, 1988):

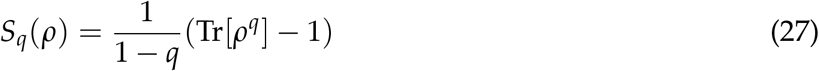

It is straightforward to generalize *S*_*w*,2_, *S*_*b*,2_ and *S*_*f*,2_ to an arbitrary *q*, and the expected Tsallis entropy depends only on 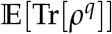. For *q* > 2 and *q* being an integer, this expectation should involve co-ancestry probabilities with more “co-ancestry arrows” and more individuals than those shown in Fig. 1B, and it perhaps contains information on higher-order correlation. Since Tsallis entropy is useful for nonextensive statistics, this general form may be of use in quantifying more complex correlation structures of local ancestry signals.

## 4. Materials and Methods

### 4.1 SLiM simulations of neutral admixture

The neutral admixture model was simulated using the standard Wright-Fisher neutral model in SLiM-4.0.1 (Haller and Messer, 2023). Admixture was not modeled explicitly, rather, we randomly assigned ancestry to ancestors according to admixture fraction **p**. Then, local ancestry in extant samples was observed from the tree sequences along their genomes that were recorded in SLiM (Haller et al., 2019). For each population size *N*, five independent realizations of the W-F model were stored as tree sequences files at each specified generation. To calculate the average entropy of a chromosomal block of length *L*, we took all non-overlapping blocks of length *L* and averaged their entropies, then averaged over several rounds of re-assignments of ancestry at the first generation, and finally averaged over the five independent realizations.

### 4.2 Numerical representation of quantum states and fast entropy calculation

Each pure state |*k*〉 is represented as a unit column vector with the *k*-th element being one. Thus, for ancestry signal *i*, its numerical form is

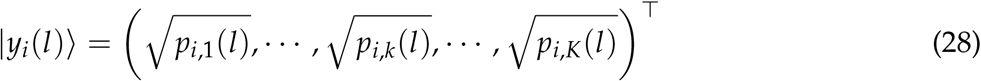

where *p_i,k_*(*l*) is the contribution at position *l* from the *k*-th source in signal *i*. The calculation of *S*_*b*,2_ follows the definition exactly, which is to first calculate the correlation kernel matrix **B** of size *n* × *n*, and then take the sum of squares of its entries. The complexity of this algorithm scales with 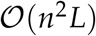. However, calculating *S*_*w*,2_ based on its definition will result in an algorithmic complexity of 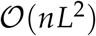, which is too large since *L* ≫ 1. Instead, we adopt the following faster but equivalent approach. First, concatenate all ancestry signals into a large array of size *nK*:

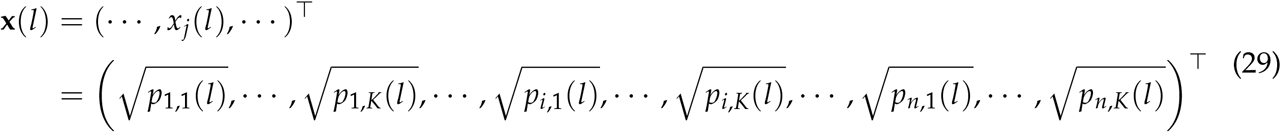

Next, calculate the correlation between components of **x**(*l*):

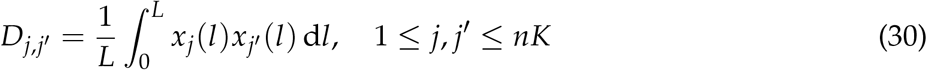

*D_j,j’_* forms an *nK* × *nK* dimensional matrix which shares the same eigenspectrum as *A*. The complexity of this approach is only 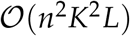. All other quantities can be calculated using their definitions.

### 4.3 Re-analysis of butterfly data

A corrected reference genome of *Papilio bianor* (Xiong et al., 2022b) was used to re-map reads from those butterfly samples in Xiong et al. (2022a) with the same processing pipeline. Then, local ancestry was estimated using ELAI (Guan, 2014) on four diploid males from population WN, while treating populations KM and XY as pure species (ELAI generation parameter = 5000). Since SNP density is high, we used the simple average among SNPs to approximate the positional average over 0 ≤ *l* ≤ *L*.

## 5. Data Availability

Scripts used in the analysis are at https://github.com/tzxiong/2023_Entropy_Ancestry_Tracts

## 6. Author Contributions

T.X. and K.B. contributed equally to the design, analysis, and writing of theoretical results. T.X. performed numerical experiments and analyzed sequence data from the butterfly hybrid zone.

## 7. Acknowledgement

T.X. is supported by a student teaching fellowship at Harvard University, and a departmental faculty fund awarded to James Mallet at Harvard University. K.B. is supported by ARO Grant W911NF-19-1-0302 and ARO MURI Grant W911NF-20-1-0082. We thank Castedo Ellerman for insightful discussion about entropy and population genetics.

## Footnotes

1 Another approach is to remove the orthogonality assumption between pure states and let their inner product be the sequence-level similarity of parental populations, so that undifferentiated parents naturally lead to hybrid offspring with highly similar homologous haplotypes. However, this modification changes the concept of ancestry and invalidates our probabilistic interpretation of entropy.

2 Note that this inequality shares a similar structure to the well-known Heisenberg uncertainty principle in physics.

