## Supplementary Information for "Quantum entropy reveals chromosomal disorder of ancestry tracts in genetic admixture"

#### 1 Expected entropy

In this section, we derive a probabilistic interpretation of the expected linear entropy in admixture from $K$ sources with proportions $\mathbf{p} = (p_1, \cdots, p_k, \cdots, p_K)$. Admixture is instantaneous at time zero, and $n$ haplotypes are sampled at time $t$.

**Definition 1** (Notation). *Throughout the text, we use the overline symbol to denote averaging a function over all combinations of positional arguments. For instance,*

$$\begin{aligned}\overline{f(l_1)} &= \frac{1}{L}\int_0^L f(l_1)\,\mathrm{d}l_1 \\ \overline{f(l_1,l_2)} &= \frac{1}{L^2}\int_0^L\int_0^L f(l_1,l_2)\,\mathrm{d}l_1\,\mathrm{d}l_2 \\ \overline{f(l_1,l_2,l_3)} &= \frac{1}{L^3}\int_0^L\int_0^L\int_0^L f(l_1,l_2,l_3)\,\mathrm{d}l_1\,\mathrm{d}l_2\,\mathrm{d}l_3\end{aligned} \tag{S1}$$

**Theorem 1.** *Using the notation from the main text, let $Q_b(l_1,l_2)$ be the probability that two different haplotypes share ancestry at both loci $l_1$ and $l_2$, and let $Q_w(l_1,l_2)$ be the probability that two loci $l_1$ and $l_2$ share ancestry in both haplotypes. Further, let $q_w(l_1,l_2)$ be the probability that two loci share ancestry in a single haplotype. The expected linear entropy is*

$$\begin{aligned}\mathbb{E}[S_{b,2}] &= 1 - \frac{n-1}{n}\overline{Q_b} - \frac{1}{n} \\ \mathbb{E}[S_{w,2}] &= 1 - \frac{n-1}{n}\overline{Q_w} - \frac{1}{n}\overline{q_w}\end{aligned} \tag{S2}$$

*Proof.* The expected $S_{b,2}$ is

$$\mathbb{E}[S(\mathbf{B})] = 1 - \frac{1}{n^2}\mathbb{E}[\mathrm{Tr}(\mathbf{B}^2)] = 1 - \frac{1}{n^2}\sum_i\sum_j \mathbb{E}[b_{ij}^2] \tag{S3}$$

For each element $b_{ij}$,

$$\mathbb{E}[b_{ij}^2] = \mathbb{E}\left[\left(\frac{1}{L}\int_0^L \langle y_i(l)|y_j(l)\rangle\,\mathrm{d}l\right)^2\right] = \frac{1}{L^2}\int_0^L\int_0^L \mathbb{E}[\langle y_i(l_1)|y_j(l_1)\rangle\langle y_i(l_2)|y_j(l_2)\rangle]\,\mathrm{d}l_1\,\mathrm{d}l_2 \tag{S4}$$

Similarly, the expected $S_{w,2}$ is

$$\mathbb{E}[S(A)] = 1 - \frac{1}{L^2}\mathbb{E}[\mathrm{Tr}(A^2)] = 1 - \frac{1}{L^2}\int_0^L \int_0^L \mathbb{E}[A^2(l_1, l_2)]\,\mathrm{d}l_1\,\mathrm{d}l_2 \tag{S5}$$

For each element $A(l_1, l_2)$,

$$\mathbb{E}[A^2(l_1, l_2)] = \mathbb{E}\left[\left(\frac{1}{n}\sum_i \langle y_i(l_1)|y_i(l_2)\rangle\right)^2\right] = \frac{1}{n^2}\sum_i\sum_j \mathbb{E}[\langle y_i(l_1)|y_i(l_2)\rangle\langle y_j(l_1)|y_j(l_2)\rangle] \tag{S6}$$

The problem narrows down to evaluate

$$\mathbb{E}[\langle y_i(l_1)|y_j(l_1)\rangle\langle y_i(l_2)|y_j(l_2)\rangle] \quad \text{and} \quad \mathbb{E}[\langle y_i(l_1)|y_i(l_2)\rangle\langle y_j(l_1)|y_j(l_2)\rangle] \tag{S7}$$

Since we assume that samples are phased to $n$ haplotypes, if $i \neq j$,

$$\begin{aligned}
&\mathbb{E}[\langle y_i(l_1)|y_j(l_1)\rangle\langle y_i(l_2)|y_j(l_2)\rangle] = \mathbb{P}(\text{Haplotypes } i \text{ and } j \text{ share ancestry at both loci } l_1 \text{ and } l_2) \\
&= Q_b(l_1, l_2) \\
&\mathbb{E}[\langle y_i(l_1)|y_i(l_2)\rangle\langle y_j(l_1)|y_j(l_2)\rangle] = \mathbb{P}(\text{Loci } l_1 \text{ and } l_2 \text{ share ancestry in both haplotypes } i \text{ and } j) \\
&= Q_w(l_1, l_2)
\end{aligned} \tag{S8}$$

If $i = j$,

$$\begin{aligned}
&\mathbb{E}[\langle y_i(l_1)|y_i(l_1)\rangle\langle y_i(l_2)|y_i(l_2)\rangle] = 1 \\
&\mathbb{E}[\langle y_i(l_1)|y_i(l_2)\rangle\langle y_i(l_1)|y_i(l_2)\rangle] = \mathbb{P}(\text{Loci } l_1 \text{ and } l_2 \text{ share ancestry in haplotype } i) \\
&= q_w(l_1, l_2)
\end{aligned} \tag{S9}$$

Since there are $n^2 - n$ pairs of $(i, j)$ with $i \neq j$ and $n$ pairs of $(i, i)$, substituting these probabilities into previous equations yields the result. □

**Theorem 2.** *Let $q_b(l_1, l_2)$ be the probability that locus $l_1$ in a randomly sampled haplotype shares ancestry with locus $l_2$ in another randomly sampled haplotype. The expected entropy on admixture fraction is*

$$\mathbb{E}[S_{f,2}] = 1 - \frac{n-1}{n}\overline{q_b} - \frac{1}{n}\overline{q_w} \tag{S10}$$

*Proof.* Recall that admixture fraction over a collection of ancestry signals is the fraction of local ancestry projected onto each pure state. Thus, the entropy of these fractions is

$$S_{f,2} = 1 - \sum_k \left[\frac{1}{nL}\sum_{i=1}^{n}\int_0^L \mathbb{I}_{|k\rangle}(|y_i(l)\rangle)\,\mathrm{d}l\right]^2 \tag{S11}$$

where $\mathbb{I}$ is the identify function. Taking expectation on both sides and expand the squares,

$$\begin{aligned}\mathbb{E}[S_{f,2}] &= 1 - \frac{1}{n^2L^2}\int_0^L\int_0^L\sum_k\left[\sum_{i=1}^n\sum_{j=1}^n\mathbb{E}[\mathbb{I}_{|k\rangle}(|y_i(l_1)\rangle)\mathbb{I}_{|k\rangle}(|y_j(l_2)\rangle)]\right]\mathrm{d}l_1\,\mathrm{d}l_2 \\ &= 1 - \frac{1}{n^2L^2}\sum_{i=1}^n\sum_{j=1}^n\int_0^L\int_0^L\left[\sum_k\mathbb{E}[\mathbb{I}_{|k\rangle}(|y_i(l_1)\rangle)\mathbb{I}_{|k\rangle}(|y_j(l_2)\rangle)]\right]\mathrm{d}l_1\,\mathrm{d}l_2\end{aligned} \tag{S12}$$

Note that $\sum_k\mathbb{E}[\mathbb{I}_{|k\rangle}(|y_i(l_1)\rangle)\mathbb{I}_{|k\rangle}(|y_j(l_2)\rangle)]$ is equal to $q_b(l_1,l_2)$ if $i\neq j$, and it is equal to $q_w(l_1,l_2)$ if $i=j$. Thus, the expression is equivalent to

$$\mathbb{E}[S_{f,2}] = 1 - \frac{n-1}{n}\overline{q_b} - \frac{1}{n}\overline{q_w} \tag{S13}$$

□

**Theorem 3.** *Take two distinct samples of haplotypes, and index them with i and j, respectively. We interpret $Q_b(l_1,l_2)$ as the probability that a random haplotype from the first sample and another random haplotype from the second sample share ancestry at both loci $l_1$ and $l_2$. Then*

$$\mathbb{E}[S_{c,2}] = 1 - \overline{Q_b} \tag{S14}$$

*Proof.* Recall that

$$S_{c,2} = 1 - \frac{1}{n_1n_2}\mathrm{Tr}[\mathbf{C}^\dagger\mathbf{C}] \tag{S15}$$

where $\mathbf{C}$ contains the correlation $c_{ij}$ of ancestry signals on each pair of haplotypes between two samples. Since

$$\mathrm{Tr}[\mathbf{C}^\dagger\mathbf{C}] = \sum_j\sum_i c_{ij}^\dagger c_{ij} = \sum_j\sum_i c_{ij}^2 \tag{S16}$$

We have

$$\mathbb{E}[S_{c,2}] = 1 - \frac{1}{n_1n_2}\sum_j\sum_i\mathbb{E}[c_{ij}^2] = 1 - \mathbb{E}[c_{ij}^2] \tag{S17}$$

Also,

$$\mathbb{E}[c_{ij}^2] = \frac{1}{L^2}\int_0^L\int_0^L\mathbb{E}[\langle y_{2,j}(l_1)|y_{1,i}(l_1)\rangle\langle y_{2,j}(l_2)|y_{1,i}(l_2)\rangle]\,\mathrm{d}l_1\,\mathrm{d}l_2 = \overline{Q_b} \tag{S18}$$

This completes the proof. □

### 2 Entropy in neutral admixture

#### 2.1 Ancestral recombination graph

Probabilities $Q_b(l_1,l_2)$, $Q_w(l_1,l_2)$, $q_b(l1,l2)$, and $q_w(l_1,l_2)$ can be evaluated using ancestral recombination graphs (ARG) embedded in a haploid population of size $2N$. Rather than running the ARG to the last coalescent event, we need to stop ARG at the time of admixture and calculate these probabilities. For $Q_b(l_1, l_2)$ and $Q_w(l_1, l_2)$, since they are for a pair of haplotypes, there are in total 15 states of the system depending on the linkage between loci and whether homologous alleles coalesce (Fig. S1). For $q_w(l_1, l_2)$, there is only one haplotype in the sample, and the stochastic process can be viewed as the previous ARG starting from state 14. For $q_w(l_1, l_2)$, it is equivalent to starting the ARG from state 15.

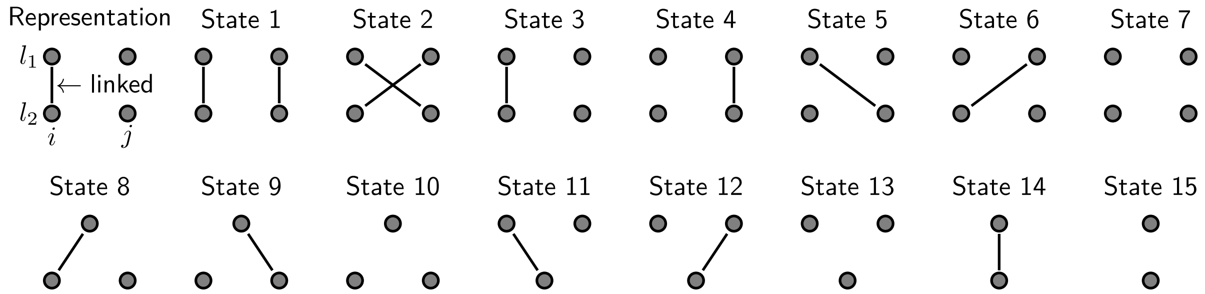


**Figure S1:** Ancestral recombination graphs with two haplotypes, two loci have 15 distinct states. Each dot is a distinct allele, and two alleles connected by solid lines are physically linked. State 1 is the initial state of the system, representing two haplotypes with two loci. States 14 and 15 represent two long-term states where homologous alleles have coalesced.

Let $\rho = 2Nr$, the generator $\mathbf{G}$ of this continuous-time process is

$$
\frac{1}{2N}\begin{pmatrix}
0 & 0 & 1 & 1 & 0 & 0 & 0 & 0 & 0 & 0 & 0 & 0 & 0 & 0 & 0 \\
0 & 0 & 0 & 0 & 1 & 1 & 0 & 0 & 0 & 0 & 0 & 0 & 0 & 0 & 0 \\
\rho & 0 & 0 & 0 & 0 & 0 & 1 & 0 & 0 & 0 & 0 & 0 & 0 & 0 & 0 \\
\rho & 0 & 0 & 0 & 0 & 0 & 1 & 0 & 0 & 0 & 0 & 0 & 0 & 0 & 0 \\
0 & \rho & 0 & 0 & 0 & 0 & 1 & 0 & 0 & 0 & 0 & 0 & 0 & 0 & 0 \\
0 & \rho & 0 & 0 & 0 & 0 & 1 & 0 & 0 & 0 & 0 & 0 & 0 & 0 & 0 \\
0 & 0 & \rho & \rho & \rho & \rho & 0 & 0 & 0 & 0 & 0 & 0 & 0 & 0 & 0 \\
0 & 0 & 1 & 0 & 0 & 1 & 0 & 0 & 0 & 1 & 0 & 0 & 0 & 0 & 0 \\
0 & 0 & 0 & 1 & 1 & 0 & 0 & 0 & 0 & 1 & 0 & 0 & 0 & 0 & 0 \\
0 & 0 & 0 & 0 & 0 & 0 & 1 & \rho & \rho & 0 & 0 & 0 & 0 & 0 & 0 \\
0 & 0 & 1 & 0 & 1 & 0 & 0 & 0 & 0 & 0 & 0 & 0 & 1 & 0 & 0 \\
0 & 0 & 0 & 1 & 0 & 1 & 0 & 0 & 0 & 0 & 0 & 0 & 1 & 0 & 0 \\
0 & 0 & 0 & 0 & 0 & 0 & 1 & 0 & 0 & 0 & \rho & \rho & 0 & 0 & 0 \\
1 & 1 & 0 & 0 & 0 & 0 & 0 & 1 & 1 & 0 & 1 & 1 & 0 & 0 & 1 \\
0 & 0 & 0 & 0 & 0 & 0 & 0 & 0 & 0 & 1 & 0 & 0 & 1 & \rho & 0
\end{pmatrix}
+ \frac{1}{2N}\,\mathrm{diag}\{-2\rho-1, -2\rho-1, -\rho-3, -\rho-3, -\rho-3, -\rho-3, -6, -\rho-1, -\rho-1, -3, -\rho-1, -\rho-1, -3, -\rho, -1\} \tag{S19}
$$

Conditioning on the ARG stopping at a specific state $m$ ($1 \le m \le 15$) at the time of admixture, one can calculate the two conditional probabilities of co-ancestry:

$$
\begin{aligned}
\mathbb{E}[\langle y_i(l_1)|y_j(l_1)\rangle\langle y_i(l_2)|y_j(l_2)\rangle|m] &= \alpha_{1,m} \\
\mathbb{E}[\langle y_i(l_1)|y_i(l_2)\rangle\langle y_j(l_1)|y_j(l_2)\rangle|m] &= \alpha_{2,m}
\end{aligned}
\tag{S20}
$$

All such probabilities are given by

$$
\boldsymbol{\alpha}_1 = \begin{pmatrix} 1-H_0 \\ 1-H_0 \\ 1-J_0 \\ 1-J_0 \\ 1-J_0 \\ 1-J_0 \\ (1-H_0)^2 \\ 1-H_0 \\ 1-H_0 \\ 1-H_0 \\ 1-H_0 \\ 1-H_0 \\ 1-H_0 \\ 1 \\ 1 \end{pmatrix}, \quad \boldsymbol{\alpha}_2 = \begin{pmatrix} 1 \\ 1-H_0 \\ 1-H_0 \\ 1-H_0 \\ 1-J_0 \\ 1-J_0 \\ (1-H_0)^2 \\ 1-H_0 \\ 1-H_0 \\ 1-J_0 \\ 1-H_0 \\ 1-H_0 \\ 1-J_0 \\ 1 \\ 1-H_0 \end{pmatrix}
\tag{S21}
$$

where $H_0 = 1 - \sum_k p_k^2$ is the probability that samples of size two comes from different ancestries at the starting point of admixture, and $J_0 = 1 - \sum_k p_k^3$ is defined similarly for samples of size three. Using these symbols, the co-ancestry probabilities in Eq. S8 for haplotypes sampled at time $t$ can be expressed as

$$
\begin{aligned}
Q_b(l_1, l_2) &= \mathbf{e}_1^\top \exp(t\mathbf{G}^\top)\boldsymbol{\alpha}_1 \\
Q_w(l_1, l_2) &= \mathbf{e}_1^\top \exp(t\mathbf{G}^\top)\boldsymbol{\alpha}_2 \\
q_w(l_1, l_2) &= \mathbf{e}_{14}^\top \exp(t\mathbf{G}^\top)\boldsymbol{\alpha}_2
\end{aligned}
\tag{S22}
$$

where $\mathbf{e}_m = (0, \cdots, 1, \cdots, 0)^\top$ is a unit vector with the $m$-th element being one.

**Theorem 4** (Short-term behavior)**.** *The initial change of expected linear entropy at the onset of admixture is:*

$$
\begin{aligned}
\partial_t \mathbb{E}[S_{b,2}(t)]|_{t=0} &= \frac{n-1}{n}\left[2(J_0 - H_0)\bar{r} - \frac{1}{2N}H_0\right] \\
\partial_t \mathbb{E}[S_{w,2}(t)]|_{t=0} &= \frac{2n-1}{n}H_0\bar{r}
\end{aligned}
\tag{S23}
$$

*where*

$$
\bar{r} = \overline{r(l_1, l_2)} \tag{S24}
$$

*is the average pairwise recombination probability in the chromosomal block* $[0, L]$.

*Proof.* Using the probabilistic interpretation Eq. S2, we have

$$\begin{aligned}\partial_t \mathbb{E}[S_{b,2}(t)] &= -\frac{n-1}{n}\partial_t \overline{Q_b}(t) \\ \partial_t \mathbb{E}[S_{w,2}(t)] &= -\frac{n-1}{n}\partial_t \overline{Q_w}(t) - \frac{1}{n}\partial_t \overline{q_w}(t)\end{aligned} \tag{S25}$$

At time $t = 0$,

$$\begin{aligned}\partial_t Q_b(l_1, l_2)|_{t=0} &= \mathbf{e}_1^\top \mathbf{G}^\top \boldsymbol{\alpha}_1 = \left(\frac{1}{2N} + 2r\right) H_0 - 2rJ_0 \\ \partial_t Q_w(l_1, l_2)|_{t=0} &= \mathbf{e}_1^\top \mathbf{G}^\top \boldsymbol{\alpha}_2 = -2rH_0 \\ \partial_t q_w(l_1, l_2)|_{t=0} &= \mathbf{e}_{14}^\top \mathbf{G}^\top \boldsymbol{\alpha}_2 = -rH_0\end{aligned} \tag{S26}$$

Averaging these quantities over positional arguments involves only averaging recombination probabilities $r = r(l_1, l_2)$. Thus, the initial change of expected entropy is

$$\begin{aligned}\partial_t \mathbb{E}[S_{b,2}(t)]|_{t=0} &= -\frac{n-1}{n}\overline{\partial_t Q_b(l_1, l_2)|_{t=0}} = \frac{n-1}{n}\left[2\bar{r}(J_0 - H_0) - \frac{1}{2N}H_0\right] \\ \partial_t \mathbb{E}[S_{w,2}(t)]|_{t=0} &= -\frac{n-1}{n}\overline{\partial_t Q_w(l_1, l_2)|_{t=0}} - \frac{1}{n}\overline{\partial_t q_w(l_1, l_2)|_{t=0}} = \frac{2n-1}{n}\bar{r}H_0\end{aligned} \tag{S27}$$

□

**Theorem 5** (Deterministic long-term behavior). *In an infinitely large population, the long-term limit of linear entropy is*

$$\begin{aligned}\lim_{t\to+\infty} \mathbb{E}[S_{b,2}(t)|N = \infty] &= 1 - \frac{n-1}{n}(1 - H_0)^2 - \frac{1}{n} \\ \lim_{t\to+\infty} \mathbb{E}[S_{w,2}(t)|N = \infty] &= 1 - \frac{n-1}{n}(1 - H_0)^2 - \frac{1}{n}(1 - H_0)\end{aligned} \tag{S28}$$

*Proof.* For each pair of loci, let $N \to \infty$ in the generator of ARG, then the three co-ancestry probabilities are:

$$\begin{aligned}Q_b(l_1, l_2) &= 2\left[(2 - H_0) H_0 - J_0\right] e^{-rt} - \left[(3 - H_0) H_0 - 2J_0\right] e^{-2rt} + (1 - H_0)^2 \\ Q_w(l_1, l_2) &= \left[H_0 e^{-rt} + (1 - H_0)\right]^2 \\ q_w(l_1, l_2) &= H_0\left(e^{-rt} - 1\right) + 1\end{aligned} \tag{S29}$$

At $t \to +\infty$, $Q_b = Q_w = (1 - H_0)^2$, and $q_w = 1 - H_0$, independent of positions. This yields the result. □

**Theorem 6** (Stochastic long-term behavior). *When genetic drift exists, the long-term limit of en-*

*tropy becomes smaller:*

$$\lim_{t\to+\infty} \mathbb{E}[S_{b,2}(t)] = 0$$
$$\lim_{t\to+\infty} \mathbb{E}[S_{w,2}(t)] = \overline{2Nr/(1+2Nr)H_0} \tag{S30}$$

*Proof.* In a finite population, genetic drift will eventually purge all haplotype diversity, and so $Q_b(l_1, l_2) \equiv 1$. This means that the long-term limit of $S_{b,2}$ is zero. For $Q_w(l_1, l_2)$, since all haplotypes are the same, $Q_w(l_1, l_2) = q_w(l_1, l_2)$. For $q_w(l_1, l_2)$, we need to solve the ARG between only state 14 and state 15. This produces

$$q_w(l_1, l_2) = \frac{2Nr\left(e^{-\frac{t}{2N}-rt} - 1\right)}{2Nr+1} H_0 + 1 \tag{S31}$$

At $t \to +\infty$, $q_w(l_1, l_2) = 1 - 2NrH_0/(2Nr+1)$. Substituting into the entropy formula for $S_{w,2}$ gives the final result. Note that if $2Nr \gg 1$ for most pairs of loci within a chromosomal block, the positional average $\overline{2Nr/(1+2Nr)}$ is close to one, and the long-term limit of $\mathbb{E}[S_{w,2}]$ is approximately $H_0$. □

#### 2.2 The entropy on admixture fractions

To calculate the entropy associated with admixture fractions among all samples using ARG, we need to evaluate an additional co-ancestry probability $q_b(l_1, l_2)$, which is the probability that position $l_1$ in a haplotype shares ancestry with position $l_2$ in a different haplotype. With the same ARG defined before, this is equivalent to starting the ARG from state 15 (two unlinked positions) and evaluate co-ancestry probability:

$$q_b(l_1, l_2) = \mathbf{e}_{15}^{\top} \exp(t\mathbf{G}^{\top})\alpha_2 \tag{S32}$$

#### 2.3 Irreversible entropy change and multilocus ancestry clusters

In the main text we define $\Delta S$ to be the difference between $S_{b,2}$ and $S_{w,2}$. The sign of this quantity reflects whether ancestry is more correlated between haplotypes ($< 0$) or within haplotypes ($> 0$). Here we prove some results on the behavior of $\Delta S$.

**Theorem 7** (Irreversible change due to recombination). *In an infinitely large population, recombination on average decreases $\Delta S$:*

$$\partial_t \mathbb{E}[\Delta S | N = \infty] < 0 \tag{S33}$$

*Proof.* Using the deterministic solutions of the three co-ancestry probabilities in Eq. S29, we have

$$\partial_t \mathbb{E}[\Delta S|N=\infty] = \frac{1}{n} r \left\{ 2e^{-2rt} \left[ J_0(2n-2) - H_0(3n-3) \right] + e^{-rt} \left[ H_0(2n-3) - J_0(2n-2) \right] \right\} \tag{S34}$$

Because $n \geq 1$ and $0 < H_0 \leq J_0$, the second term in the bracket is negative:

$$H_0(2n-3) - J_0(2n-2) = (2n-2)(H_0 - J_0) - H_0 < 0 \tag{S35}$$

For the first term in the bracket,

$$J_0(2n-2) - H_0(3n-3) = (n-1)(2J_0 - 3H_0) = (n-1)\left(3\sum_k p_k^2 - 2\sum_k p_k^3 - 1\right) \tag{S36}$$

It remains to be shown that $3\sum_k p_k^2 - 2\sum_k p_k^3 - 1 \leq 0$. We use Lagrange multiplier to find the maximum of the augmented function

$$F(p_1, \cdots, p_K) = 3\sum_k p_k^2 - 2\sum_k p_k^3 - 1 + \lambda(\sum_k p_k - 1) \tag{S37}$$

Local extrema of the augmented function satisfy

$$\begin{cases} \partial_{p_k} F = 6p_k - 6p_k^2 + \lambda = 0 \\ \partial_\lambda F = \sum_k p_k - 1 = 0 \end{cases} \tag{S38}$$

The first equation has solutions

$$\begin{cases} p_{k,-} = \frac{1}{2} - \frac{1}{6}\sqrt{9+6\lambda} \\ p_{k,+} = \frac{1}{2} + \frac{1}{6}\sqrt{9+6\lambda} \end{cases} \tag{S39}$$

If $9 + 6\lambda > 0$, any two $p_k$s cannot be both $p_{k,+}$ in order to keep the sum smaller than one. If one is $p_{k,+}$ and another is $p_{k,-}$, then all other $p_k$s are zero, thus zero must be a solution and $p_{k,-} = 0$. This also means that $F = 0$. If all $p_k$s are $p_{k,-}$, the distribution is at maximum entropy, and $p_k = 1/K$, thus $F = 3/K - 1/K^2 - 1 \leq 0$. Finally, if $9 + 6\lambda = 0$, then both $p_k$s are $1/2$, and $F$ is also nonpositive. Thus, $F \leq 0$ holds for all discrete distributions.

Consequently, the entire bracket in Eq. S34 is negative for all possible $H_0$ and $J_0$, which completes the proof. □

**Conjecture 1** (Irreversible change under recombination and genetic drift). *The general ARG of neutral admixture has*

$$\partial_t \mathbb{E}[\Delta S] < 0 \tag{S40}$$

Since

$$\partial_t \mathbb{E}[\Delta S] = \frac{n-1}{n}\partial_t(\overline{Q_w} - \overline{Q_b}) + \frac{1}{n}\partial_t \overline{q_w} \tag{S41}$$

And $\partial_t \overline{q_w} < 0$, it suffices to show that $\partial_t(\overline{Q_w} - \overline{Q_b}) \leq 0$. It is then sufficient to show that for each pair of loci $l_1$ and $l_2$, $\partial_t Q_w(l_1, l_2) - \partial_t Q_b(l_1, l_2) \leq 0$, which brings us back to the two-haplotype, two-locus ARG. In Fig. S2 and S3, numerical evaluation of $Q_w(l_1, l_2) - Q_b(l_1, l_2)$ under various parameters shows that it is indeed a decreasing function through time.

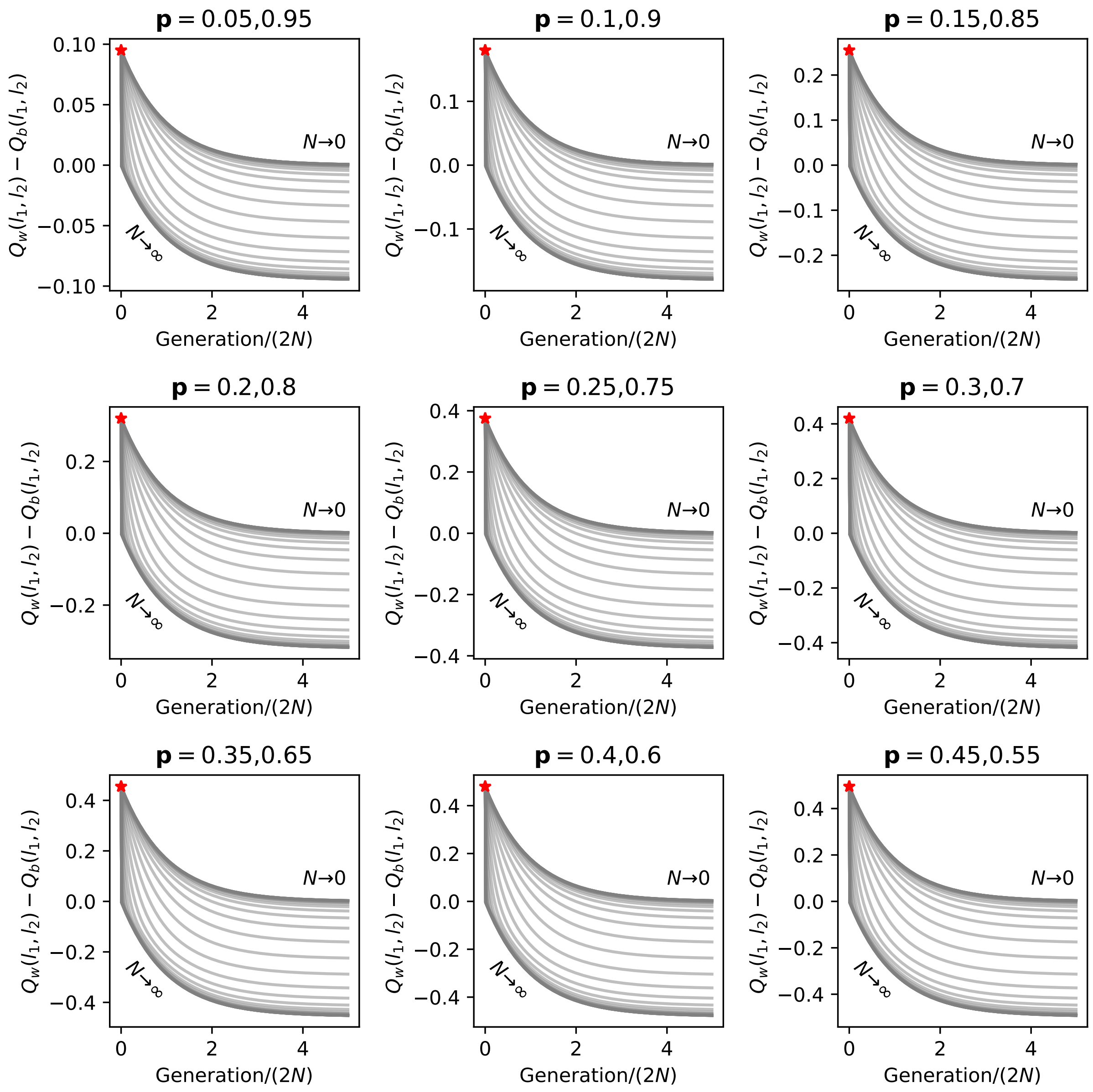


**Figure S2:** Change in the difference between two co-ancestry probabilities through time under the full ARG. Admixture is from two sources with proportions **p**. Values are numerically computed using the ARG generator. Red stars represent the starting state at $t = 0$. Each trajectory represents a fixed $\rho = 2Nr$. The range of $\rho$ is $10^{-5}$ to $10^{5}$. Time is measured in generation scaled by $2N$.

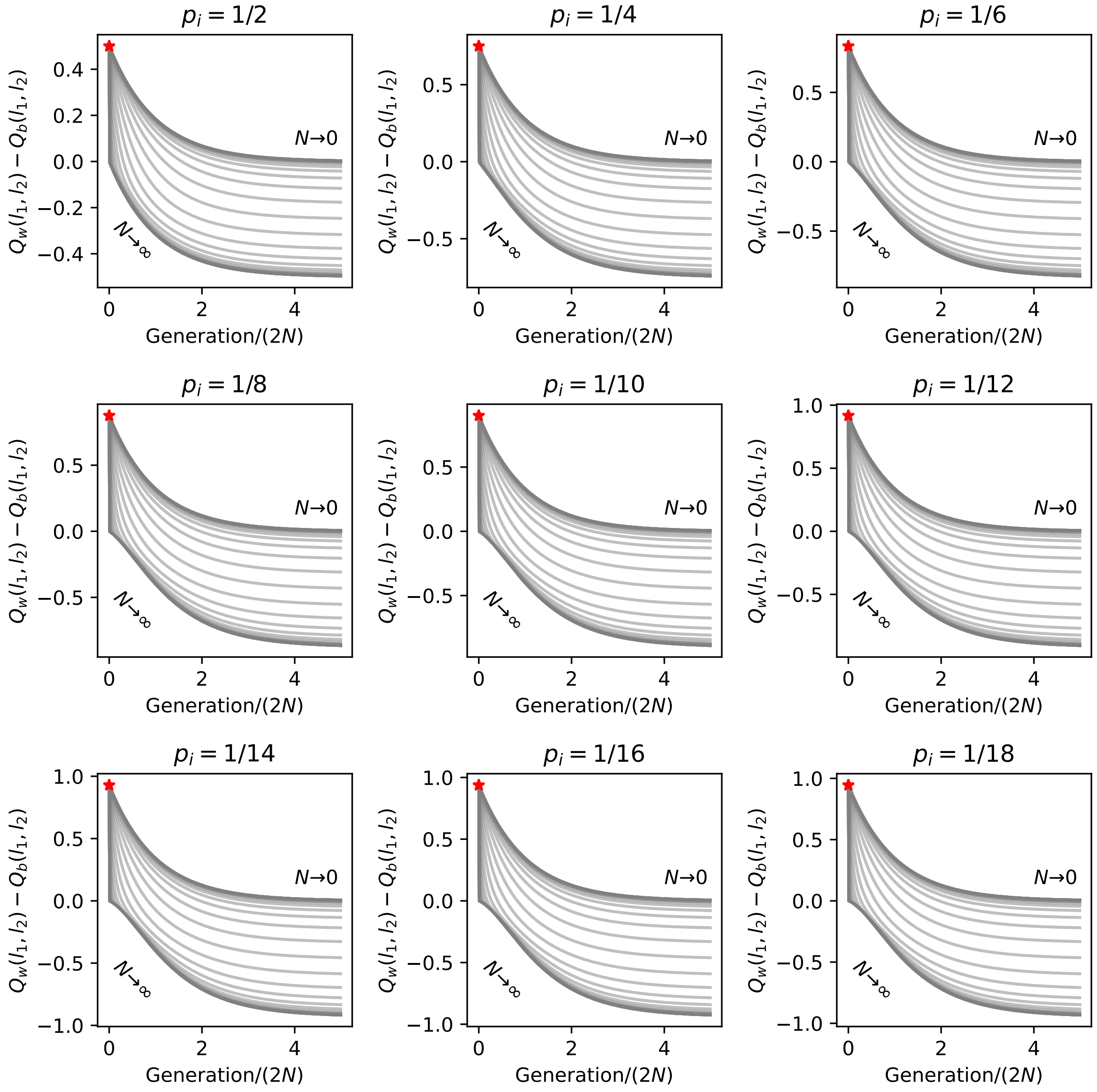


**Figure S3:** Change in the difference between two co-ancestry probabilities through time under the full ARG. Admixture is from $K$ sources with proportion $p_i = 1/K$ for all $i$. Values are numerically computed using the ARG generator. Red stars represent the starting state at $t = 0$. Each trajectory represents a fixed $\rho = 2Nr$. The range of $\rho$ is $10^{-5}$ to $10^{5}$. Time is measured in generation scaled by $2N$.

### 3 Re-analysis of butterfly hybrid genomes

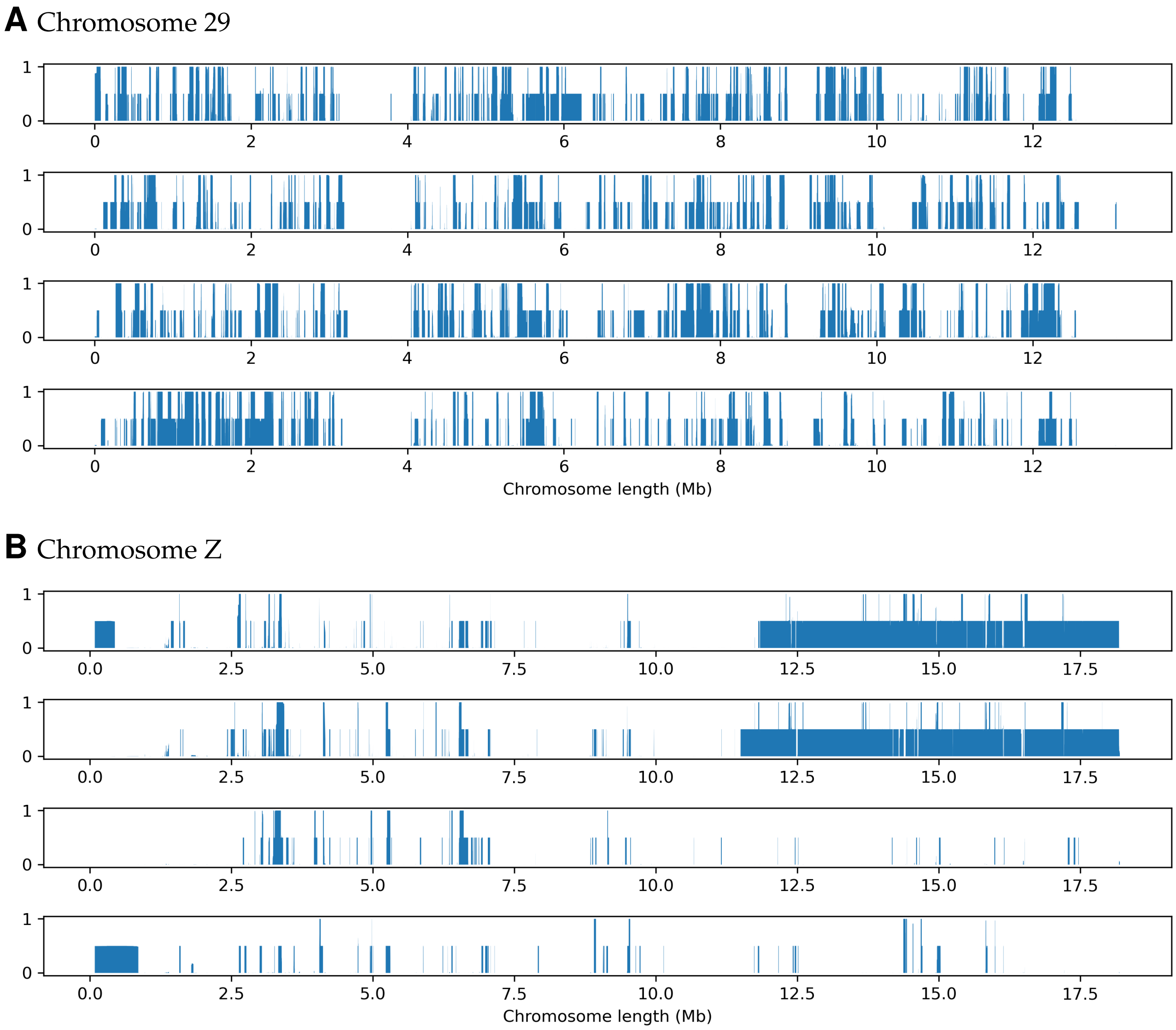


**Figure S4:** Local ancestry estimated by ELAI in four hybrid individuals on two chromosomes. The Z chromosome resembles either pure species or those with early-generation backcross ancestry tracts. Blue is associated with contribution from *P. maackii*
